# LEAP: A Generalization of the Landau-Vishkin Algorithm with Custom Gap Penalties

**DOI:** 10.1101/133157

**Authors:** Hongyi Xin, Jeremie Kim, Sunny Nahar, Can Alkan, Onur Mutlu

## Abstract

**Motivation:** Approximate String Matching is a pivotal problem in the field of computer science. It serves as an integral component for many string algorithms, most notably, DNA read mapping and alignment. The improved LV algorithm proposes an improved dynamic programming strategy over the banded Smith-Waterman algorithm but suffers from support of a limited selection of scoring schemes. In this paper, we propose the Leaping Toad problem, a generalization of the approximate string matching problem, as well as LEAP, a generalization of the Landau-Vishkin’s algorithm that solves the Leaping Toad problem under a broader selection of scoring schemes.

**Results:** We benchmarked LEAP against 3 state-of-the-art approximate string matching implementations. We show that when using a bit-vectorized de Bruijn sequence based optimization, LEAP is up to 7.4x faster than the state-of-the-art bit-vector Levenshtein distance implementation and up to 32x faster than the state-of-the-art affine-gap-penalty parallel Needleman Wunsch Implementation.

**Availability:** We provide an implementation of LEAP in C++ at github.com/CMU-SAFARI/LEAP.

## 1 Introduction

Approximate String Matching, also known as fuzzy string matching, is a classic problem in the realm of string algorithms. The goal is to match a *search* or *pattern* string to the *reference* string while allowing for errors (Navarro [2001]). There are numerous formulations to this problem, depending on the relative lengths of the pattern and the reference strings. For similarly sized strings, the problem is often finding the optimal sequence of operations (called *edits*) needed to transform the pattern string into the reference. Common edits include insertions, deletions, and substitutions, each of which have their own associated penalty score. Similarity between the pattern and the reference is then quantified by the total *penalty score* sum of all edits. The goal of approximate string matching is to find the optimal sequence of edits that minimizes the total penalty score incurred in transforming the pattern into the reference. In the case where the pattern string is much smaller than the reference, the problem often becomes identifying substrings of the reference that best match the pattern.

Approximate string matching has a wide array of applications in computer science. It is one of the core compute units in modern Next-Generation-Sequencing (NGS) short read aligners (or *mappers*), such as MUMmer (Delcher et al. [1999]), BWA (Li and Durbin [2009]), SOAP2 (Li et al. [2009]), mrFAST (Hach et al. [2010], Xin et al. [2013]), SNAP (Matei et al. [2011]), and Bowtie (Langmead and Salzberg [2012]). The purpose of sequence alignment is to find the most likely location where the (*read*) originated from a given reference. Based on the assumption that individuals of the same species have highly similar genomes, a representative *reference genome* (Flicek and Birney [2009]) is constructed for a species, and all subsequent intra-species genome sequencing heavily relies on this reference. DNA reads are matched to the reference genome to find potential mappings of the read in the target genome under a suitable error threshold. When the similarity between the read and the reference segment is high, the mapper concludes (with high probability) that the read must have been sampled from the same position in the unknown target genome. These potential locations are subsequently used to determine the final position of the reads for assembly of the genome.

For short read aligners, approximate string matching computation is a large percentage of the overall runtime. This is because of two reasons: complex genome variations and large reference genome sizes. There are tens of millions of reads which need to be matched to the reference genome, and each read needs to be checked against a large number of possible locations in the reference, depending on the mapping strategy. Therefore, developing a faster and more efficient matching algorithm, that is also highly parallel, is paramount. This becomes more important as modern compute infrastructures become increasingly more parallel and SIMD friendly.

A wide variety of algorithms have been developed to solve the approximate string matching problem and its variations (Cole and Hariharan [2002], Ukkonen [1985]). These include optimized bit-vector implementations (Myers [1999], Baeza-Yates and G. Navarro [1999]) and algorithms used for alignment (Needleman and Wunsch [1970], Smith and Waterman [1981]). While these improvements increase the performance of the algorithm through parallelization, most implementations still follow the basic dynamic-programming doctrine of filling a partial scoring matrix, which records the optimal alignment between the strings of the string pair, through a top-to-bottom, left-to-right manner.

An alternative to the canonical dynamic programming strategy of filling an *L × L* matrix, (assuming the two strings are of equal length *L*), is to iteratively find the longest matching substrings with an increasing number of edits. This concept was first proposed by Landau and Vishkin (Landau and Vishkin [1989]) and is often called the Landau-Vishkin algorithm (although also called Landau-Vishkin, this is not the Landau-Vishkin algorithm published in 1986 (Landau and Vishkin [1986])). For simplicity, we refer to the Landau-Vishkin algorithm (Landau and Vishkin [1989] as LV for the reminder of this paper. One limitation with LV is that it was only proposed for Levenshtein distance scoring schemes and it is not proven to work for more general scoring schemes.

In this paper, we present an extension to the previously proposed Landau-Vishkin (Landau and Vishkin [1989]) algorithm, which is an optimization over the Smith-Waterman algorithm specifically for banded global alignment with Levenshtein distance penalty scores. We show that the same principle of Landau-Viskhin can be applied not only to global approximate string matching with Levenshtein distance penalty scores, but also to any banded global or semi-global approximate string matching problems with non-negative scoring schemes. To achieve this, we first propose a generalization of the approximate string matching problem called the *Leaping Toad* problem and show that all banded global and semi-global approximate string matching problems with positive penalty scores can be transformed into the Leaping Toad problem. Then we propose **LEAP**, a general dynamic-programming solution for the Leaping Toad problem based on the Landau-Vishkin algorithm. Finally we provide a bit-vectorized de Bruijn sequence based optimization over LEAP. We show that LEAP is 7.4x faster than the state-of-the-art bit-vector Levenshtein distance implementations and 32x faster than the state-of-the-art parallel affine gap penalty Needleman-Wunsch implementations.

This paper makes the following contributions:

- It proposes the Leaping Toad problem, a generalization of all banded global or semi-global approximate string matching problems with positive penalty scores. It then shows the detailed procedure of transforming approximate string matching problems with Levenshtein distance penalty scores and affine-gap penalty scores into the Leaping Toad problem.
- It provides a new algorithm, LEAP, an extension of the Landau-Vishkin’s algorithm, that solves the general Leaping Toad problem.
- It provides a detailed proof of the optimality of LEAP. The proof confirms that LEAP captures the minimum-score edit sequence between the two strings, under any positive penalty scoring scheme.
- It provides a bit-vectorized, de Bruijn sequence based optimization over LEAP, which uses a perfect hash function that exploits properties of de Bruijn sequences to find the position of the most significant ‘1’ in a bit-vector with simple bit-vector operations.
- It shows that bit-vectorized LEAP is 7.4x faster than the state-of-the-art Levenshtein approximate string matching implementations and up to 32x faster than the state-of-the-art affine-gap approximate string matching implementations.

The rest of the paper is organized as follows; the Background section provides a detailed explanation of the approximate string matching problem, as well as past work and analyses of common solutions; the Methods section describes the Leaping Toad problem, as well as its general solution, LEAP, and the bit-vector optimization over LEAP; the Results section compares LEAP against the state-of-the-art Levenshtein distance implementations as well as affine-gap implementations; the Discussion section discusses the advantages and limitations of LEAP; and finally, the Conclusion section summarizes the paper.

## 2 Background

The Landau-Vishkin (Landau and Vishkin [1989]) algorithm improves upon the banded-edit distance algorithm. It uses the fact that edit-distance is conserved along the diagonal for a sequence of matches, so it can simply traverse along the diagonal to the position of the next error. The length of the traversal is `LCE`(*i, j*) (longest common extension), which is the length of the longest prefix which *s*_*i..m*_ and *r*_*j..n*_ share.

The Landau-Vishkin algorithm uses a variant matrix for edit-distance computation: `LV`_*d,e*_ stores the maximal row along diagonal *d* with edit distance *e*, where *d* is calculated as *j - i*, where *i* is the row and *j* is column in the edit-distance matrix. By conditioning on the last error, the recurrence for `LV` follows:

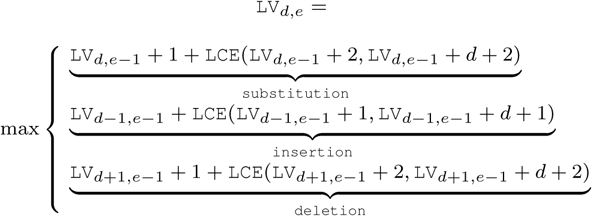

## 3 Methods

Both the global and semi-global Banded Levenshtein Distance Problem (BLDP) and Banded Affine Gap Distance Problem (BAGDP) can be generalized as a restricted optimal path finding problem in a directed acyclic graph. We call this the *Leaping Toad* problem (LTP). In this section, we first propose the Leaping Toad problem, and we show how a general edit-distance problem can be converted to an instance of LTP. Subsequently we propose an improved dynamic programming algorithm **LEAP** as a solution, followed by a proof of its optimality. We discuss the backtracking process of LEAP. In addition, we provide a de Bruijn sequence based bit-vector optimization over LEAP. Finally, we discuss specific optimizations to the algorithm for affine-gap penalties.

**Fig. 1.**
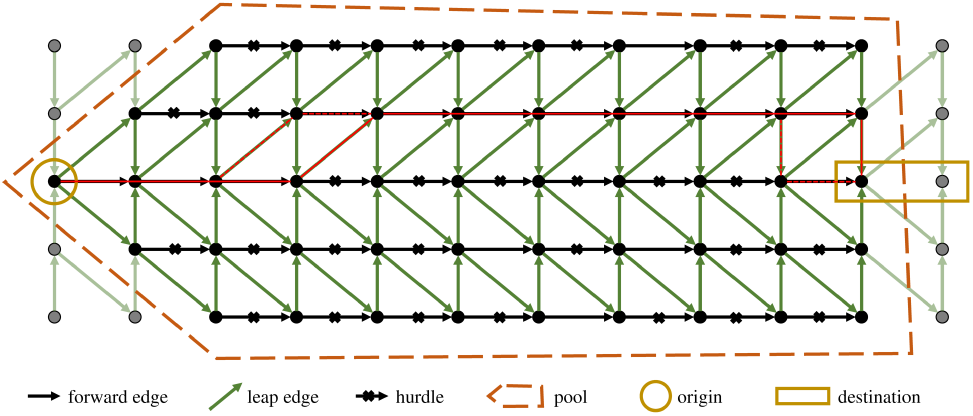
This figure shows a swimming pool setup that is equivalent to the banded Levenshtein distance problem. In this setup, black edges are forward edges. Black crosses are hurdles on the edge. Green edges are leap edges. All leaps and hurdles cost +1 energy. The goal is to find an optimal path from the origin vertex to the destinations while consuming minimum energy. Both the solid and dashed red lines are legitimate optimal paths for this swimming pool setup.

### 3.1 The Leaping Toad Problem

The Leaping Toad Problem (or simply **LTP**) can be summarized as a traversal problem in a special directed acyclic graph, where a toad travels in the weighted graph and the goal is to find a path that connects the origin and the destination vertices while minimizing the sum of edge costs along the path.

The directed acyclic graph of LTP is described as follows:

- There is a **convex** *swimming pool* that encircles vertices taken from a 2-dimensional vertex grid, where vertices are aligned in rows and columns. Vertices in the swimming pool are then organized into disjoint *lanes* which are rows in the vertex grid. Inside a lane, each vertex is connected to the next vertex on its right by a directional edge, with itself being the source and the vertex on the right being the destination. We call these edges as *forward edges*. Forward edges only exist among vertices inside the swimming pool and do not exist for vertices outside of the swimming pool.
- A vertex may also have edges pointing to vertices in other lanes. We call these edges *leap edges*. In LTP, for a vertex and a separate lane, there can be at most one leap edge pointing to at most one vertex in that lane. In other words, there can not be multiple edges pointing to the same lane from the same vertex. We also require all the vertices in the same lane share the same types of leap edges: the same directions and lengths. When visualized, leap edges between two lanes are an array of parallel arrows. Notice that some leap edges might have their source and/or destination vertices staying out of the swimming pool and we call these edges *out edges*. Outside of the swimming pool enclosure, out edges continue to exist, connecting vertices between different lanes.
- Over any edge, there is a **non-negative integer** weight. Leap edges sharing the same origin and destination lanes have the same, **positive** weight. Forward edges have zero or positive weights. We call forward edges with positive weights as *hurdles*. Traveling across a hurdle is called *hurdle crossing*. Hurdles may have different costs.

In the swimming pool, we appoint a number of lanes as *origin lanes* and a number of lanes as *destination lanes*. The set of origin lanes and destination lanes may overlap. The general **goal** of the LTP is to find a path in the directed graph, with minimum sum of edge weights, that starts at the first vertex (the leftmost vertex in the swimming pool of the lane) of an origin lane and **either travels to the last vertex of a destination lane or travels out of the swimming pool while exiting onto a destination lane**.

**Fig. 2.**
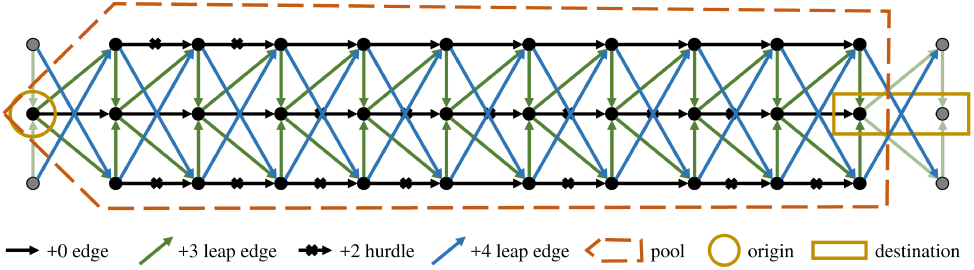
This is an alternative swimming pool setup. Compare to the pool in Figure 1. In this pool, each vertex can have leaping edges pointing to all lanes. Furthermore, in this setup, crossing a hurdle costs +2 energy, leaping a single lane costs +3 energy, and leaping two lanes costs +4 energy.

For simplicity, we call edge weights as *energy costs*; we call traveling along the leaping edge as *leaps*. Also for the simplicity of developing a solution, we require all the leaping edges to never point backwards to vertices in previous columns on the left. A relaxation of this restriction is discussed in the Discussion section.

Figure 1 shows an example setup of the swimming pool as well as the optimal path to cross the pool (in red). In this setup, as the figure shows, the toad starts at the first vertex in the middle lane on the left side of the pool and the goal is to travel to the last vertex of the middle lane on the right side of the pool. In a lane, black crosses are placed on the hurdle edges.

In this particular setup, the toad can only leap to neighboring lanes, as the arrow shows. The leaping edges are set differently depending on whether the lane is 1) the middle lane, 2) above the middle lane or 3) below the middle lane. If it is in the middle lane, leaping edges are tilted by 45 degrees pointing to the vertex in the next column as they point to neighboring lanes. For other lanes, leap edges are vertical when they point towards the center lane and are tilted by 45 degrees when they point away from the center lane. Here we also set the energy cost of hurdles as well as leaps as 1. The red line depicts an optimal path for the toad to travel across the pool. Notice that there can be multiple optimal paths with the same total energy cost (shown as dashed red lines).

There are many alternative setups to LTP. For an alternative setup, a number of settings could be changed:

1. The energy cost of overcoming different hurdles can be different.
2. There can be leap edges pointing to more lanes.
3. The energy cost of leaps can be random and lane specific.

Figure 2 shows an alternative setup.

### 3.2 Conversion of Approximate String Matching to the Leaping Toad problem

Both the banded Levenshtein and the banded affine gap string matching problems can be converted to an instance of LTP. To convert both problems into LTP, we first convert both string matching problems into an optimal path finding problem in a directed graph. Then we show that the optimal path problem in the converted directed graph is indeed an instance of LTP.

The Banded Levenshtein Distance Problem (BLDP) can be easily converted into an optimal path finding problem in a directed graph. For simplicity, in this paper we assume BLDP takes a pair of equal-length strings, such as strings *r, s* of length *L*. For the (*L* + 1) *×* (*L* + 1) edit-distance matrix *D*, we assign each element *D*_*i,j*_ of the matrix a unique vertex *v*_*i,j*_. Using the edit-distance recurrence function, a directional edge is drawn from vertex *v*_*i,j*_ to *v*_*i′*_, _*j′*_ if and only if *v*_*i,j*_ *≠ v*_*i′*_,_*j′*_ and *i*^*′*^*-i ≤* 1 and *j*^*′*^ *- j ≤* 1 (an edge to the right, bottom and bottom-right element). On each edge (*v*_*i,j*_ *, v*_*i′*_, _*j′*_), we place an integer weight *w*, where *w* = 0 if *i*^*′*^ = *i* + 1, *j*^*′*^ = *j* + 1, and *s*_*i*_ = *r*_*j*_, or *w* = 1 otherwise. An example of the directed graph representation of the Levenshtein distance problem is shown in Figure 3. The objective function of BLDP becomes an optimal path finding problem where we want to find a path with minimum total edge weight within the edit-distance threshold *e* from *v*_0,0_ to *v*_*L,L*_.

**Fig. 3.**
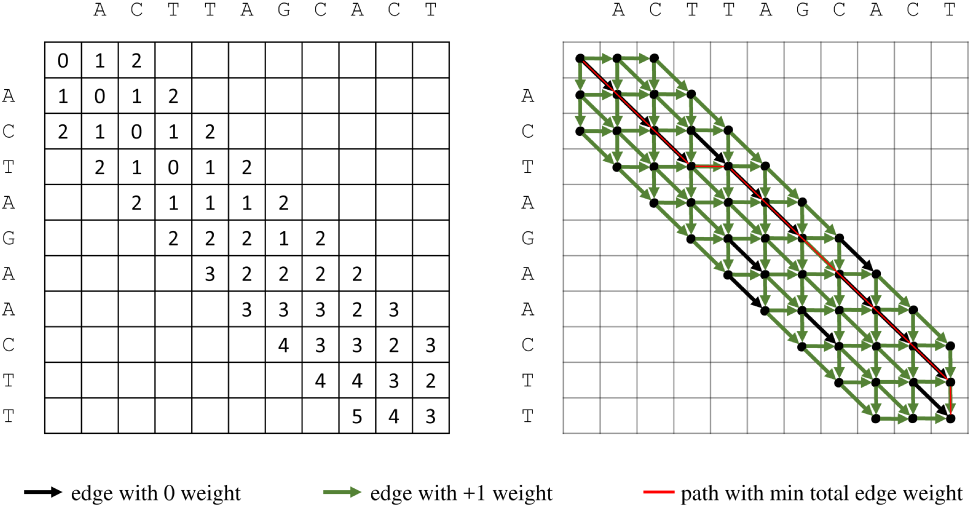
This figure shows the directed graph representation of the dynamic-programming method of the banded Levenshtein distance problem. Each vertex in the graph represents a element in the DP matrix. The red line outlines the optimal path traveling from the top-left vertex to the bottom-right vertex.

For BLDP, the equivalent swimming pool directed graph setup is shown in the example in Figure 1. We call this the Levenshtein Leaping Toad setup. In general, given a (*N* + 1) *×* (*N* + 1) LDP matrix and a maximum edit-distance threshold *e*, we formulate the equivalent Levenshtein Leaping Toad setup as the following:

The swimming pool has 2*e* + 1 lanes. The lane *l* contains *L - |l|* vertices, specifically the set of vertices *v*_*i,j*_ such that *i-j* = *l*. As Figure 1 shows, the right-end of all lanes are aligned while the left-end of the lanes forms a wedge shape. Each lane in the swimming pool corresponds to a diagonal in the edit-distance matrix by construction: the center lane represents the central diagonal and the *k*th lane above or below the center lane represents the *k*th diagonal above or below the central diagonal.

After mapping vertices and edges accordingly, we can observe that a hurdle is placed in the lane *l* between the *k*th and the *k* + 1st vertex if the corresponding edge in the BLDP graph has nonzero weight. Also for each lane in the pool, there are leap edges pointing to neighboring lanes. For the center lane (*l* = 0), leap edges are always tilted by 45 degrees. When they are on any other lane, leap edges are vertical when they are pointing towards the center lane and are tilted by 45 degrees when they are pointing away from the center lane. All hurdles and leaps cost 1 unit of energy.

After converting BLDP to the Levenshtein Leaping Toad setup, the **goal** becomes:

1. Determine if the toad can swim from the first vertex of the center lane to the last of the center lane while spending at most *E* energy.
2. If it can, then find the path that costs the minimum amount of energy. While there is slightly different from the goal of LTP, which allows traveling out of the swimming pool and allows terminating on the center lane but outside of the swimming pool, we will show later that for BLDP with small edit distance budget *E*, the result path of LTP is either the same with BLDP, or can be easily transformed into the path of BLDP. For now, we the goal of LTP is equivalent to the goal of BLDP.

The equivalence between the two directional graphs of LTP and BLDP can be visualized in Figure 4. For the vertex at the *k*th column of the *l*th lane above (or below) the center lane in the swimming pool, we assign its equivalent vertex in the Levenshtein direction as the vertex of element *E*_*x*_,_*y*_ in the edit-distance matrix *E*, where *x* = *k - l* and *y* = *k* (or *x* = *k* and *y* = *k - l*). For example, the red, green, and blue vertices highlighted in both figures are equivalent between the two graphs, as well as their inbound and outbound edges. It’s worth noting that vertices in the same column in the swimming pool forms a mirrored “L” shape in the BLDP graph (highlighted as purple lines in both graphs in the figure), with the vertex of the center lane on the corner. Also, no leap edges point backwards. All leap edges point either to vertices in the same column or in the next column.

**Fig. 4.**
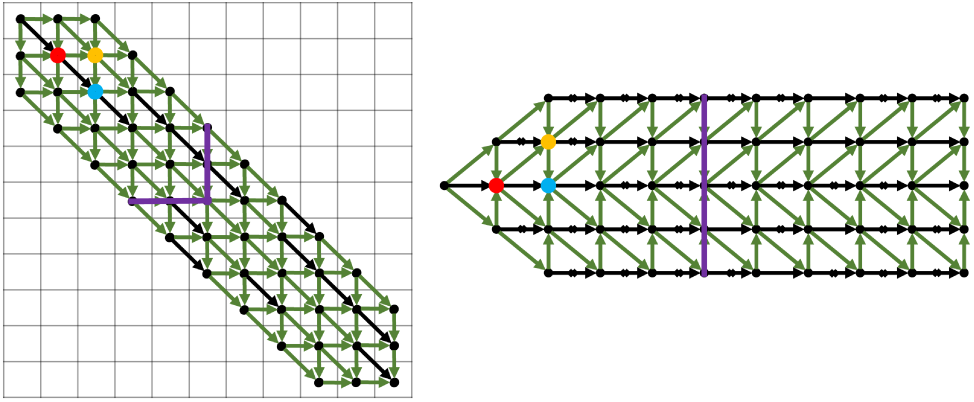
This figure shows the equivalence between the directed graph in Figure 3 and the Leaping Toad setup in Figure 1. Notice that the red vertices, the yellow vertices and the blue vertices are equivalent between the two graphs. The purple line outlines the equivalent vertices in the directed graph of an entire column of vertices of the swimming pool.

For semi-global alignment, the objective function of BLDP changes from finding an optimal path **from the top-left element of the matrix to the bottom-right element of the matrix** (global alignment), to finding an optimal path **from any element of the first column of the matrix to any element of the last column of the matrix** (semi-global alignment), we simply need to reflect the same changes in LTP. Therefore for semi-global alignment, the objective function in the Leaping Toad setup changes to finding an optimal path from **the first vertex of bottom half lanes to the destination of top half lanes** with minimum energy cost.

For Banded Affine Gap Distance Problem (BAGDP), or in general for any banded custom-gap-penalty string-matching problem (with a maximum insertion or deletion threshold *k*, a maximum energy budget *E* and positive gap penalties for different gap lengths), as now the toad can take an arbitrary length insertion or deletion (within the insertion limit *k*) and each insertion length has its own penalty score, we modify the swimming pool setup as each lane can have leap edges pointing to up to *k* lanes above and below the lane. Leap edges having the same origin but different destinations will have different energy costs. Figure 2 shows the leaping toad setup for a global BAGDP with a specific affine gap penalty scheme, where mismatches are penalized with +2, gap openings are penalized with +3 and gap extensions are penalized with +1.

### 3.3 LEAP: The general solution of the Leaping Toad Problem

Similar to approximate string matching problems, the Leaping Toad Problem can be solved through dynamic-programming. Since we restrict the toad from ever going backward, the toad can only reach a vertex from another vertex that is from the same or previous column. Therefore, for each new column, we can find the optimal paths leading to its vertices, as well as the minimum traveling energy, by reusing the optimal path results from previous columns: for each new vertex, we find all prior-vertices that can reach to the target vertex in one step, then pick the prior-vertex that requires the least amount of combined energy of both reaching to the prior-vertex and the intermediate move. We repeat this process until either we have reached a destination vertex, or no vertices in a column is still within the energy budget.

A major drawback of the above naïve dynamic-programming solution of the Leaping Toad Problem, as with the naïve dynamic-programming solution of the edit-distance problem, is that for each new vertex, we have to compare all of its previous-step vertices, then pick the vertex with the minimum overall energy cost. Similarly in backtrack, as we move one step backward at a time, we have to once again resolve the previous-step vertex for each and every vertex along the optimal path.

Inspired by the Landau-Vishkin algorithm for the edit-distance problem, we propose an improvement over the naïve solution of the Leaping Toad problem. We only consider switching lanes at vertices that are right before a hurdle. When there is no hurdle, we always let the toad swim forward; therefore avoid frequently checking possibilities of leaping from other lanes. We name this algorithm **LEAP**.

LEAP is developed upon a key observation that among all possible optimal paths with minimum energy costs, there must exist at least one optimal path that the toad **either never leaps or only leaps right before a hurdle or only leaps through out edges.**

Theorem 1. *Among all optimal paths of the Leaping Toad problem, there must exist one path in which the toad either never switches lanes or only switches right before hurdles.*

Before proving the theorem, we first define some terminology: we refer to a path from the origin vertex to the destination vertex simply as a *path*. The path may or may not have any lane switches. Whenever there is a lane switch, we call it a *leap*. Between two leaps, the toad only goes forward and we call such straight segments of the path as *segments*. We further categorize segments into two groups: segments that end with the destination vertex or a hurdle as *complete segments* and segments that do not end with such conditions as *incomplete segments*. We call the operation that extends the incomplete segment until it either reaches a destination vertex or a hurdle as *completing the segment*. Equipped with this terminology, we are now ready to prove the stepping-stone lemma of Theorem 1:

Lemma 1. *For a path with an incomplete segment S, there must exist an alternative path that shares the same moving sequence before S, while completing S into Sc and have at most the same cost.*

Proof. To prove the lemma, we need to find an alternative path that supports the claim. Assume in the original path, after *S*, the path continues with a series of leaps and segments, denoted as a *moving sequence*, [*L*_1_, *S*_1_, *L*_2_, *L*_3_, *S*_2_, …], where *S*_*i*_ is the *i*th segment after *S* while *L*_*j*_ is the *j*th leap after *S*. Note that between two leaps there can be either zero or one segment, while between two segments there has to be at least a single leap.

Assuming that *S*_*c*_ is *d* vertices longer than *S* (*|S*_*c*_*|* = *|S|* + *d*), we propose an alternative path that shares the same segments and leaps before *S*, followed by *S*_*c*_, and then continues with the same sequence of leaps [*L*_1_, …, *L*_*t*_], while skipping all the segments from *S*_1_ to *S*_*k–1*_, where 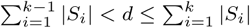, and *L*_*t*_ is right before *S*_*k*_ in the original moving sequence.

If 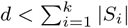, which suggests that after taking [*L*_1_, …, *L*_*t*_], the alternative path merges with the original path somewhere in *S*_*k*_, then we also add the latter half of *S*_*k*_ after the merge point. The alternative path is then completed with the same moving sequence after *S*_*k*_.

If original path does not have enough segments after *S* to match the length of *d*, then the alternative path simply takes the remaining leaps while skipping all the remaining segments. In this special case, the toad will take the out edges and leap out of the pool to finish the leaping sequence.

Compared to the original path, the alternative path is guaranteed to have at most the original energy cost. This is because:

1. The energy cost before *S* in the original path and before *Sc* in the alternative path are identical as they take identical moving sequences.
2. The energy costs of the two paths after the merge point (if they do merge) are also identical, as the two paths also take identical moving sequences.
3. The energy cost of the leaping sequence after *S*_*c*_ and before the merge point of the alternative path is at most the energy cost of the moving sequence after *S* and before the merge point in the original path. This is because the original path takes the same leaping sequence [*L*_1_,…,*L*_*t*_] (hence consumes the same amount of leaping energy) but the other segments skipped by the alternative path may contain hurdles and hence cost extra.
4. The energy costs of *S* and *S*_*c*_ are identical since *S*_*c*_ is only a completion of *S*, and by construction the extension is free of hurdles so it costs zero energy.

**Fig. 5.**
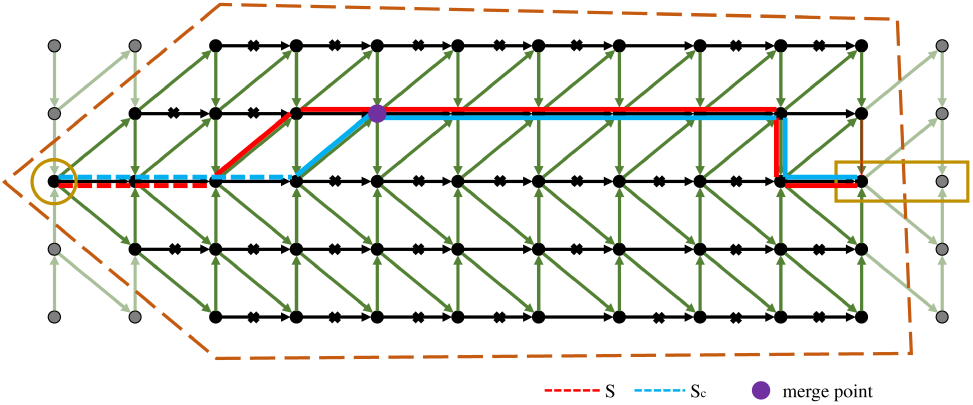
This figure illustrates the procedure of Lemma 1. In this example, we have a random optimal path in red (*ρ*_1_). We claim that by extending the first incomplete segment of the red path, *S*, into a complete segment, *S*_*c*_, we are guaranteed to find another optimal path (*ρ*_2_, in blue) with the same or smaller energy cost.

Figure 5 depicts an example of converting a segment *S* in the original path (red) into *S*_*c*_ with an alternative path (blue) using the above procedure. Compared to the red path, the blue path consumes less energy as the red path contains segments with hurdles which are skipped by the blue path. With Lemma 1, we are now ready to proof Theorem 1. We prove through contradiction:

Proof. Assume there exist no optimal paths that either never leap or only have complete segments. Then for any optimal path, it must take at least one leap and have at least one incomplete segment.

We arbitrarily pick an optimal path *ρ*_1_ and find the first incomplete segment *S*_1_ in the path. Following the procedure in Lemma 1, we can find an alternative path *ρ*_2_ without the incomplete segment *S*_1_ with the same energy cost (*ρ*_2_ cannot have a smaller energy cost; otherwise *ρ*_1_ is not optimal). *ρ*_2_ is also an optimal path, so by assumption, it must contain another incomplete segment *S*_2_. We subsequently repeat the procedure.

The above process can only iterate a finite number of times since the procedure in Lemma 1 does not introduce any new segments, as completing *S* into *S*_*c*_ maintains that segment, and the procedure skips the segment sequence [*S*_1_,…,*S*_*k-*1_] and may either shorten or skip *S*_*k*_, depending on the position of the merge point. Hence each iteration removes a incomplete segment. Given that there are finite number of incomplete segments, there can only be finite number of iterations. The final product after all iterations is a path with no incomplete segments with the same energy cost as *ρ*_1_. However, according to our assumption, such path does not exist. Hence, this leads to a contradiction, which proves Theorem 1.

With Theorem 1, we now transform the general Leaping Toad problem of *finding an optimal path with minimum cost* to a sub-problem that *finds an optimal path which only contains complete segments*. As we have proven in Theorem 1, the resulting optimal path of the sub-problem must also be an optimal path of the general Leaping Toad problem.

LEAP solves the above sub-problem through an optimized dynamic-programming method that can be viewed as an extension of the Landau-Vishkin algorithm. LEAP can be summarized into four steps:

1. LEAP iterates through all intermediate energy costs from 0 to *E* and for each energy cost, LEAP iterates through all lanes.
2. For an intermediate energy cost *e* and a lane *l*, LEAP finds the furthest vertex *v* in *l* that is reachable at precisely the energy cost *e* from either a leap or a hurdle-crossing.
3. LEAP extends the segment at *v* (if permitted) until the segment hits a hurdle.
4. LEAP repeats step 2) and 3) until either a lane has reached to the destination vertex or all intermediate energy levels have been exhausted. The path that leads to the destination vertex is reported as the result.

To summarize, LEAP uses a core recurrence function shown below:

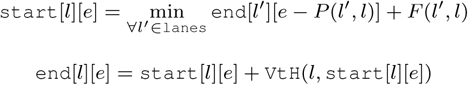

where *P* (*l′, l*) returns the penalty of leaping from lane *l′* to *l*, *F* (*l′, l*) returns the number of columns the toad moves forward when it leaps from lane *l′* to *l*, and `VtH`(*l,* `start_column`) (abbreviated for Vertices to Hurdle) returns the number of vertices until next hurdle from `start_column` in lane *l*. When *l* = *l′*, *P* (*l, l′*) is simply the energy cost of the next hurdle and *F* (*l, l′*) = 1.

The detailed pseudo-code of LEAP is shown in Algorithm 1.

Theorem 2. *The result path returned by LEAP is indeed the optimal path of the sub-problem.*

We validate the correctness of Theorem 2 using two arguments. First, all segments in the result path of LEAP is guaranteed to be complete, because in step 3, LEAP always extends a segment until it reaches a hurdle. Second, for any energy cost *e* and any lane *l*, the last vertex extended in step 3 (if any) marks the furthest vertex that the toad can reach to in *l* using precisely *e* energy. Combining both arguments, we can conclude that if LEAP can find a path that reaches to the destination vertex with energy cost *e < E* while the toad cannot reach the destination with energy cost *e′ < e*, then the energy cost of the optimal path of the Leaping Toad sub-problem must be *e* (otherwise, according to the second argument, for a smaller energy, the toad would have already reached the destination) and the result path returned by LEAP must be an optimal path.

Proof. The first argument is obvious. The second argument can be proven through induction:

*Base case*: When *e* = 0, since any leap or hurdle-crossing would consume a non-zero amount of energy, the furthest vertex the toad can reach in a lane with zero energy cost would be the last vertex in the lane before hitting a hurdle. Therefore, the second argument holds true for the base case.

*Induction step*: Assume for all intermediate energy costs *e′ < e*, and for all lanes, the second argument holds true. That is, for any lane *l′*, the last vertex `end`[*l′*][*e′*] reached by step 3 in LEAP marks the furthest vertex the toad can reach in *l′* while consuming precisely *e′* amount of energy.

Now, because both hurdle crossings and leaps cost positive amount of energy, to get to a vertex with *e ≠* 0 energy cost in lane *l*, the toad has to either leap from a vertex in another lane or cross a hurdle in the same lane from a vertex in which the the total energy cost to get to that vertex is less than *e*. Since LEAP has already calculated the furthest vertices of all lanes for all energy levels *e ′ < e* (based on our assumption), we can conclude that step 2 of LEAP will find the furthest vertex in *l* such that it is reachable from either a leap or a hurdle-crossing at precisely *e* energy cost.

Finally the only remaining method for the toad to get to a vertex while costing *e* energy, is to swim straight, without running into a hurdle, from a previous *e*-energy vertex. This vertex is also captured by LEAP in step 3. Therefore, the argument is correct for the induction step.

#### Algorithm 1: LEAP

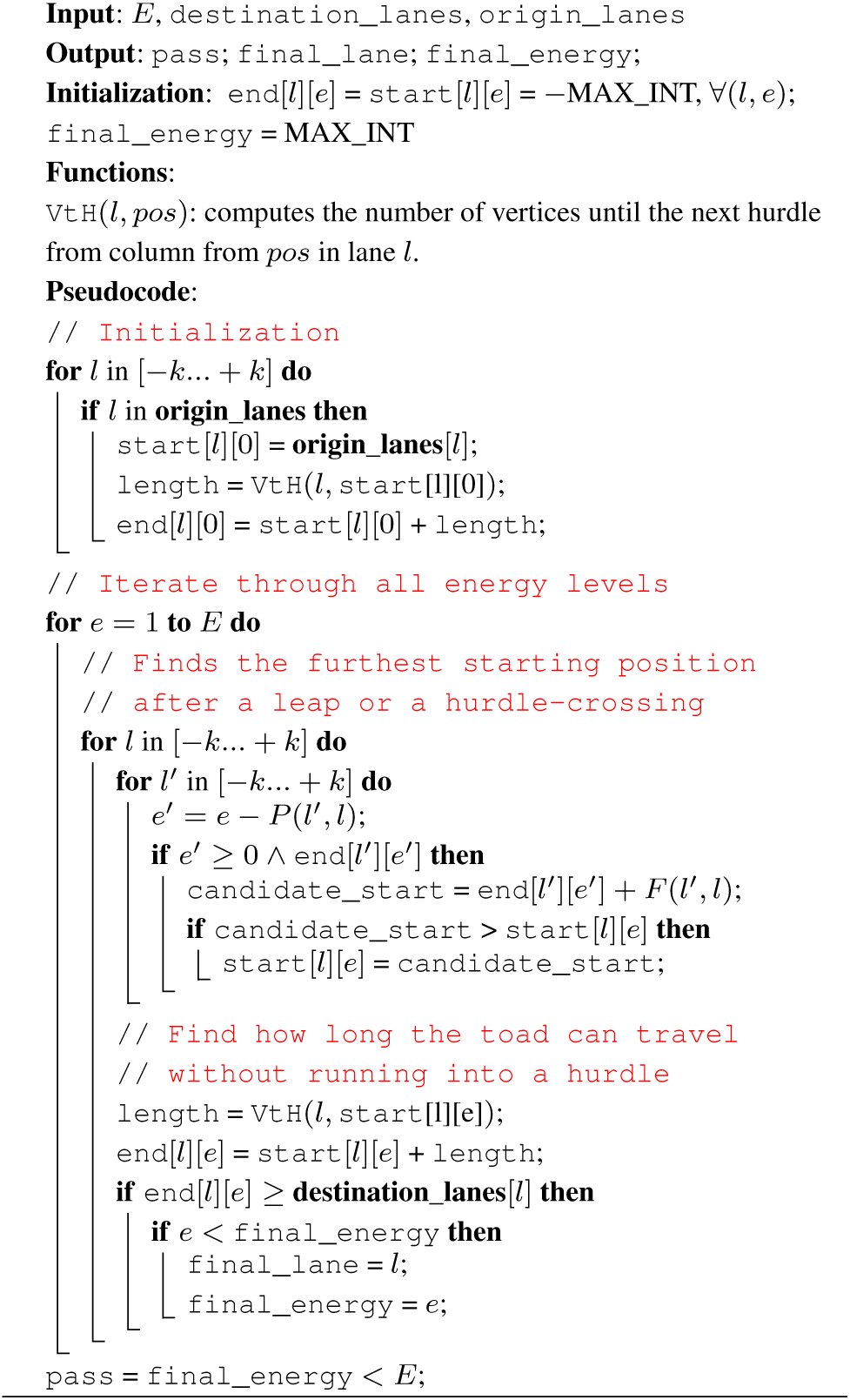

Conclusion: After step 3, LEAP always reflects the furthest vertex the to ad can reach in the target lane *l* under the target energy cost *e*.

### 3.4 Backtracking in LEAP

The pseudo-code of the backtracking method of LEAP is shown in Algorithm 2.

A remaining issue with LEAP, or LTP in general, is that it allows the toad to leap out of the swimming pool (recall the LTP and BLDP equivalent-goal assumption we made in Section 3.2). According to the LTP problem definition, as long as the toad reaches to a destination lane, even if the destination vertex is outside of the pool, the path is still acceptable. Translated back to approximate string matching problems, this sometimes leads to awkward results, since inserting or deleting letters (counterparts of leaps in the approximate string matching notion) beyond the end of the string is undefined. For instance, assume we are computing the global alignment between strings “`AAAAAC`” and “`AAAAAG`” with a simple scoring scheme: mismatches are penalized with +5, single letter gap is penalized with +4, double letter gap is penalized with +2 (this gap penalty might not make sense for DNA alignment, as here single letter gap is more costly than double letter gap). For this particular string pair, the correct edit sequence would be, `M-3I-2D-2`, which translates to, “3 matches, inserting 2 letters, then deleting 2 letters”. The minimum edit cost would be +4. When subjected to LEAP, this string pair will instead generate an edit sequence of `M-4I-2D-2`. We have `M-4` simply because LEAP does not consider taking a leap before running into a hurdle, which in approximate string matching notion, is the `C-G` mismatch. Although the energy cost of the LEAP path is still +4, the edit sequence clearly does not make sense since one cannot delete two letters after 4 matches when there is only a single letter `C` left. We can easily correct backtracking sequences from the out-of-bound LEAP backtrack sequences. Our work assumes that forward edges without hurdles always cost 0 energy. This corresponds to matches in approximate string matching and alignment. Hence, we can remove matches from the out-of-bound LEAP backtrack sequence until the length matches the intended length. This transformation maintains the total energy cost. For the above example, the intended sequence length is 7. So we have to simply remove one match, and we can remove the last `M` and transform `M-4I-2D-2` to `M-3I-2D-2`, which is an optimal edit sequence.

#### Algorithm 2: Backtrack

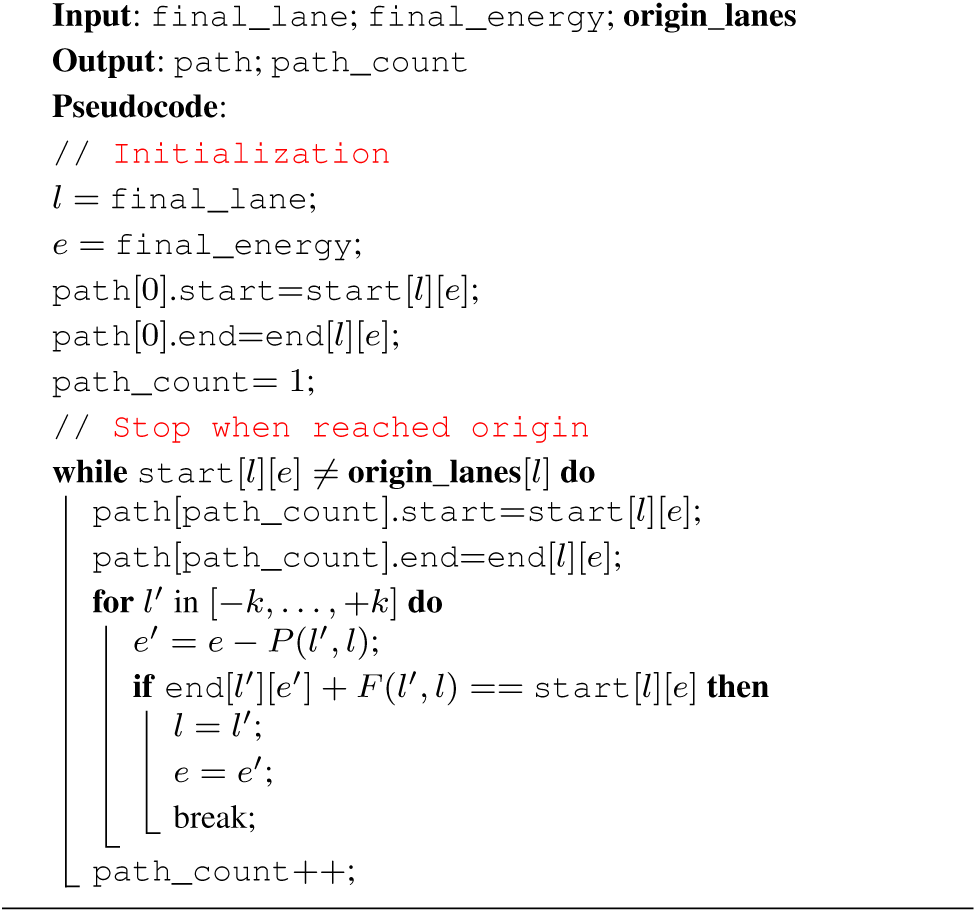

**Table 1.** The de Bruijn sequence LSB lookup table for 8 bit words.

|  |  |
| --- | --- |
| 000 | 0 |
| 001 | 1 |
| 010 | 6 |
| 011 | 2 |
| 100 | 7 |
| 101 | 5 |
| 110 | 4 |
| 011 | 3 |

### 3.5 De Bruijn Sequence Optimization

While LEAP can drastically reduce the number of comparisons in the dynamic-programming solution of the Leaping Toad problem, step 3 of LEAP still involves a costly loop that searches for the next hurdle.

By encoding the hurdle information as bit-vectors, we can significantly improve the performance of step 3 using de Bruijn sequences and bit-vector operations. The detailed technique is described in Leiserson et al. [1998]. Here we provide a brief summary of the technique.

First, we encode the sequence of all forward edges (edges that go straight) of a lane as a bit-vector, where ‘0’ denotes an edge that does not have a hurdle in between the two connected vertices while ‘1’ denotes an edge that does. For example, the middle lane in Figure 1 can be represented as ‘0001111110’.

Counting the number of edges before the next hurdle after the *i*th vertex is equivalent to counting the number of 0’s from the *i*th bit until we hit a 1. After shifting the bit-vector *i* bits to the left, the problem then becomes finding the position of the most significant 1 in the resulting bit-vector, which is equivalent to counting the number of trailing 0’s of the reverse bit-vector.

First proposed in the paper of Leiserson et al. [1998], counting the number of trailing 0’s in a bit-vector can be carried out through a hash-table lookup with a perfect hash function. Assume the machine word has a length of 2^*n*^ bits. The least significant 1 of a vector *b* can be singled out by *b* ANDed with its two’s complement number 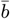 (computed through `NOT`(*b*) + 1). For example for a machine of word size of 2^3^ = 8 bits, the least significant 1 of a vector *b* = 01001000 can be singled out by *b* `AND` 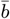 which is *b*_LSB_ = 01001000 `AND` (10110111 + 1) = 01001000 `AND` 10111000 = 00001000 (LSB stands for least significant bit). Then the number of trailing 0’s can be computed by multiplying *b*_LSB_ with a pre-computed de Bruijn sequence dB_seq_ of 2^*n*^ bits (in our example, *n* = 3 and subsequently dB_seq_ = 00011101). Because *b*_LSB_ must be power of two, *b*_LSB_ *×* dB_seq_ essentially translates to shifting *b*_LSB_ to the left *m* times, where *m* is the number of trailing zeros. By taking the most significant *n* bits of the product (carried out through shifting the product to the right 2^*n*^ *-n* bits), we have then produced a unique number, a `key`, between [0, …, 2^*n*^ *-* 1]. Finally, we can use the `key` to query a pre-computed lookup table of 2^*n*^ entries, which returns the pre-computed number of trailing 0’s in *b*_LSB_. The example lookup table for dB_seq_ = 00011101 is provided below in Table 1. The pseudo code of finding the next hurdle is shown in Algorithm 3.

#### Algorithm 3: Vertices To Hurdle

**Input:** *l*; `start_pos`

**Output:** `V_num`

**Internal Variable: bit_vec**: hurdle encoded binary bit-vectors;

*dB*_*seq*_: de Bruijn sequence of 2^*n*^ bits

**Functions:** `reverse_bits(bitvec): reversing the bit-vector sequence`

`lookup(key):` lookup the precomputed de Bruijn sequence table

**Pseudocode:**

`// Initialization`

shift_bit_vec = **bit_vec**[*l*] ≪ start_pos;

`rev_bit_vec = reverse_bits(shift_bit_vec);`

`b_LSB = rev_bit_vec ^(:(rev_bit_vec) + 1);`

key = (b_LSB × dB_seq_) ≫ 2^*n*^ − *n*;

`V_num = lookup(key)`

### 3.6 LEAP variant for affine gap penalty

Similar to the Needleman-Wunsch, LEAP can also be modified to more efficiently support affine gap penalties using separate insertion *I* and deletion *D* matrices. Instead of tracking leaps from all possible lanes, with affine gap penalty we maintain an *I* and a *D* matrix separately for each lane to track the furthest column the toad can reach to from a leap, under different energy costs. Specifically, *I*[*l*][*e*] and *D*[*l*][*e*] stores the furthest column that the toad can arrive to, from an upward or downward leap, respectively, while consuming precisely *e* energy.

With *I* and *D* arrays, we modify the core recurrence function as follows:

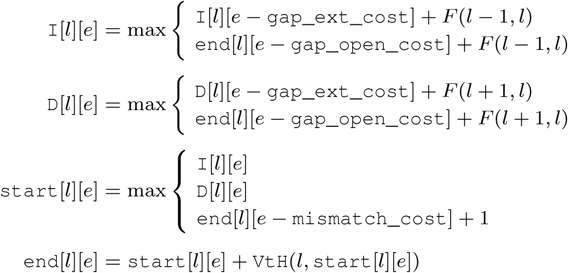

where *F* (*l-*1*, l*) = 0 if lane *l* is above the center lane and *F* (*l-*1*, l*) = 1 otherwise, and *F* (*l* + 1*, l*) = 0 if lane *l* is below the center lane and *F* (*l -* 1*, l*) = 1 otherwise.

## 4 Results

We implemented LEAP for both banded Levenshtein distance and banded affine gap penalties. For each scoring scheme, we compare LEAP against three state-of-the-art approximate string matching implementations, including: an in-house vanilla Landau-Vishkin implementation (LV); an implementation of Gene Myer’s bit-vector algorithm from SeqAn (SeqAn) (Döring et al. [2008]) and finally a SIMD implementation of banded global Needleman-Wunsch algorithm (NW-SIMD) (Daily [2016]). Additionally, in order to benchmark the benefit of the de Brujin sequence based bit-vector optimization, we implemented two versions of LEAP: one with (LEAP-BV) and one without (LEAP), the bit-vector optimization.

To benchmark the performance of the above implementations, we augmented a popular aligner, bowtie2, to dump all read and reference pairs into a separate file as ASCII string pairs, during the mapping procedure. For comprehensiveness, we gathered reads from six read files from the 1000 Genomes Project (1000 Genomes Project Consortium [2010]), ERR240726_1, ERR240726_2, ERR240727_1, ERR240727_2, ERR240728_1, ERR240728_1. Each read file is mapped against the human reference genome version 37 with bowtie2 under default settings. All reads in the above read files are 100-bp long. For banded Levenshtein distance, we benchmarked all six implementations with different edit-distance thresholds *E* ranging from 1 to 5. For banded affine gap penalties, we set the matching score as 0; the mismatch penalty as +2; gap open penalty as +3 and gap extend penalty as +1. We set the total affine gap penalty threshold to be 3 *× E*.

Finally, we conducted two separate tests. In the first test, shown in Table 2, we benchmarked all read and reference pairs from bowtie2 on all 6 implementations. In the second test, shown in Table 3, we only selected read-reference pairs that have at most five edits. While the first test evaluates the performance of different implementations under a realistic mapper environment, the second test evaluates how fast can each implementation find the optimal alignment in a highly similar string pair.

From both tables, we can observe that LEAP-BV is the fastest in both Levenshtein distance setup and affine gap setup. For Levenshtein distance, compared to SeqAn, LEAP-BV achieves up to **7.4x** speedup under *E* = 1 and **1.6x** speedup under *E* = 5. For affine gap, compared to NW-SIMD, LEAP-BV achieves even greater performance, with up to **32x** speedup under *E* = 1 and **2.3x** speedup under *E* = 5. Notice that even though both vanilla LV and SeqAn are reasonably fast under Levenshtein distance settings, neither support affine gap penalties due to their tight coupling with Levenshtein distance scores.

Furthermore, we observe that the performance of LV and LEAP are very similar. This is expected since under Levenshtein scores, LEAP reduces to LV. We also observe that the performance of both LEAP and LEAP-BV decreases with increasing *E*. This is also expected, since under a greater *E*, LEAP checks more lanes and iterates through more energy levels.

**Table 2.** Mixed DNA String Pairs (seconds / 10 million pairs). This table shows runtime for a suite of Approximate String Matching algorithms normalized to seconds per 10 million read/reference pairs. String pairs in this benchmark are generated by bowtie2 with default parameters. While LEAP uses a simple for loop to find the next hurdle, LEAP-BV (LEAP-Bit-Vector) uses the de Bruijn sequence based bit-vector algorithm to locate the next hurdle.

|  | e | LEAP-BV | LEAP | LV | NW-SIMD | SeqAn |
| --- | --- | --- | --- | --- | --- | --- |
| Levenshtein | 1 | 0.84 | 1.28 | 1.25 | 38.81 | 6.21 |
|  | 2 | 1.48 | 2.07 | 2.03 | 38.91 | 6.66 |
|  | 3 | 2.39 | 3.14 | 3.11 | 38.61 | 6.28 |
|  | 4 | 3.35 | 4.50 | 4.45 | 38.38 | 6.59 |
|  | 5 | 4.60 | 6.01 | 6.04 | 38.46 | 6.88 |
| Affine Gap | 1 | 1.18 | 1.72 | N/A | 38.75 | N/A |
|  | 2 | 3.10 | 4.01 | N/A | 38.15 | N/A |
|  | 3 | 6.39 | 7.77 | N/A | 38.64 | N/A |
|  | 4 | 10.91 | 13.31 | N/A | 38.66 | N/A |
|  | 5 | 16.91 | 20.74 | N/A | 38.51 | N/A |

**Table 3.**
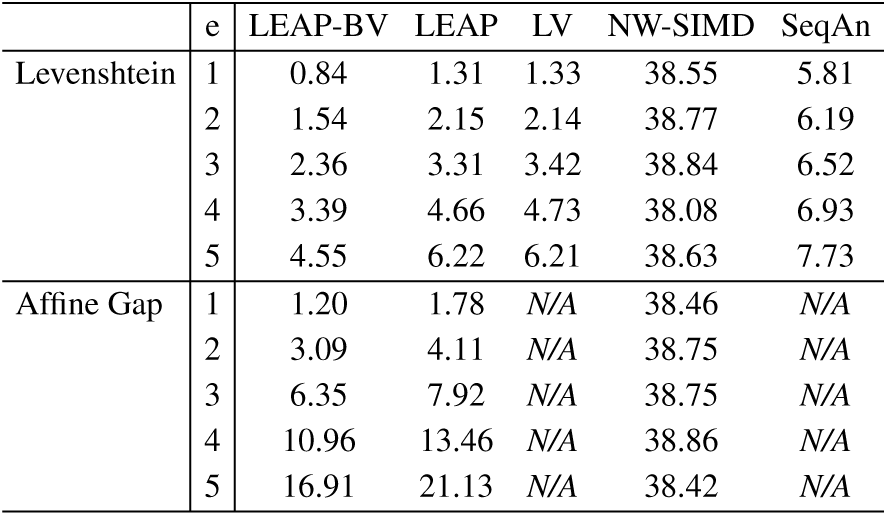
Highly Similar DNA String Pairs (seconds / 10 million pairs). This table shows runtime for a suite of Approximate String Matching algorithms normalized to seconds per 10 million read/reference pairs. String pairs in this benchmark are a subset of the ones in Table 2. String pairs in this set have at most 5 edits in total between the two strings.

Last but not the least, compared to LEAP, LEAP-BV provides on average 39% improvement.

Overall, LEAP performs best under small edit-distance thresholds, while its performance quickly decreases as the edit-distance threshold increases. Nonetheless, from our experiments, LEAP-BV is still faster than other implementations even under moderate edit-distances.

## 5 Discussion

While we required the toad to never move backwards while it leaps in the original definition of the Leaping Toad Problem, this requirement is not a necessity for LEAP. Both Theorem 1 and Theorem 2 hold true even if the toad is allowed to move backward as it leaps. The key premise in proving both theorems is that all leaps and hurdles cost positive amounts of energy while moving forward without running into a hurdle costs zero energy.

However, when the toad is allowed to move backward during a leap, the naïve column-by-column dynamic programming method stops working, since a toad could leap from a “future column” that has not yet been calculated. LEAP, on the other hand, remains intact and functional under such conditions. This makes LEAP a broader solution for a more general Leaping Toad problem compared to the naïve dynamic-programming method.

A major limitation of LEAP, in spite of its advantages, is that it cannot handle both negative energy bonuses and positive penalty schemes. In terms of approximate string matching, this translates to not supporting negative scores for matches along with positive penalties for mismatches and gaps (or vice versa). It only supports positive penalties for mismatches and gaps with no penalty/bonus for matches. As a result, LEAP cannot handle local alignment.

Nonetheless, LEAP shows great potential to be composed with NGS mappers where seed-and-extend methods are often used and strings are often compared with global or semi-global alignment.

Overall, LEAP provides three major benefits:

- LEAP reduces the frequency of calling the recurrence function, from *𝒪* (*E × N*) times to *𝒪* (*E*^2^) times (for the Levenshtein edit-distance case).
- LEAP incorporates a de Brujin sequence based hash-table optimization, which further speeds up the computation of the Leaping Toad problem.
- LEAP enables greater parallelization in solving global and semi-global alignment problems.

Unlike traditional Needleman-Wunsch and Smith-Waterman parallel implementations, which focus on exploiting parallelism between elements on the same anti-diagonal line in the dynamic programming matrix, LEAP enables a more efficient parallelization approach. As we have discussed in the de Bruijn sequence optimization subsection, LEAP utilizes hurdle-encoded bit-vectors to calculate the position of the next hurdle. In the realm of Approximate String Matching, the bit-vector of each lane is simply the letter-wise XOR between the two strings after shifts. For the center lane, the bit-vector is indeed the letter-wise XOR between the pattern and the reference string; while for the *i*th lane above (or below) the center lane is the same letter-wise XOR but after shifting the pattern (or the reference) to the right *i* times. Given that we can further encode each letter with log_2_(*σ*) bits, the entire operation of preparing all bit-vectors can be done in 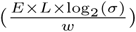 XORs, where *w* is the length of the machine word in bits. Under the *parallel random-access machine model (PRAM)*, all the XORs can be calculated in parallel for lanes under the same energy budget *e*. Lanes with different energy budgets however share dependency. Nonetheless, since the outer loop only iterates up to *E* times, (compared to the parallel Smith-Waterman or Needleman-Wunsch, whose outer loop is often iterated *L* times) LEAP still can provide a greater parallel speedup under the PRAM model.

## 6 Conclusion

Approximate string matching is an important and widely studied problem, and its use in critical components for a large number of applications has created a need to develop faster, efficient, and highly parallelizable solutions.

In this paper, we analyzed existing approximate matching algorithms such as the Smith-Waterman and Needleman-Wunsch algorithms. We reviewed the Landau-Vishkin algorithm, an fast method for calculating Levenshtein distance. We then proposed the Leaping Toad problem, a generalization of the approximate string matching problem, as well as LEAP, a generalization of the Landau-Vishkin algorithm that solves the Leaping Toad problem under a broader selection of scoring schemes. We provided a detailed proof that LEAP solves the Leaping Toad problem.

We compared LEAP against 3 state-of-the-art approximate string matching implementations. We showed that when using a bit-vectorized de Bruijn sequence based optimization, LEAP achieved a 7.4x speedup over the state-of-the-art bit-vector Levenshtein distance implementation and was up to 32x faster than the state-of-the-art affine-gap-penalty parallel Needleman Wunsch Implementation.

## 7 Acknowledgements

We thank Carl Kingsford for his valuable input into this work.

## References

1000 Genomes Project Consortium. A map of human genome variation from population-scale sequencing. Nature, 467:1061–1073, 2010.

Baeza-Yates and R. G. Navarro. Faster approximate string matching. Algorithmica, 23(2):127–158, 1999.

R. Cole and R. Hariharan. Approximate string matching: A simpler faster algorithm. SIAM Journal on Computing, 31(6):1761–1782, 2002.

J. Daily. Parasail: SIMD C library for global, semi-global, and local pairwise sequence alignments. BMC Bioinformatics, 17:81, 2016. doi: 10.1186/s12859-016-0930-z. URL http://dx.doi.org/10.1186/s12859-016-0930-z.

A. L. Delcher, S. Kasif, R. D. Fleischmann, J. Peterson, O. White, and S. L. Salzberg. Alignment of whole genomes. Nucl. Acids Res., 1999.

A. Döring, D. Weese, T. Rausch, and K. Reinert. Seqan an efficient, generic c++ library for sequence analysis. BMC Bioinformatics, 9:11, 2008. doi: 10.1186/1471-2105-9-11. URL http://dx.doi.org/10.1186/1471-2105-9-11.

P. Flicek and E. Birney. Sense from sequence reads: methods for alignment and assembly. Nature methods, 6(11 Suppl):S6–S12, Nov. 2009. ISSN 1548-7105. doi: 10.1038/nmeth.1376. URL http://dx.doi.org/10.1038/nmeth.1376.

Hach, F. Hormozdiari, C. Alkan, F. Hormozdiari, I. Birol, E. E. Eichler, and S. C. Sahinalp. mrsfast: a cache-oblivious algorithm for short-read mapping. Nature methods, 7(8):576–577, 2010.

M. Landau and U. Vishkin. Introducing efficient parallelism into approximate string matching and a new serial algorithm. In Proceedings of the Eighteenth Annual ACM Symposium on Theory of Computing, STOC ’86, pages 220–230, New York, NY, USA, 1986. ACM. ISBN 0-89791-193-8. doi: 10.1145/12130.12152. URL http://doi.acm.org/10.1145/12130.12152.

G. M. Landau and U. Vishkin. Fast parallel and serial approximate string matching. Journal of algorithms, 10(2):157–169, 1989.

Langmead and S. L. Salzberg. Fast gapped-read alignment with bowtie 2. Nature Method, 9:357–359, 2012.

E. Leiserson, H. Prokop, and K. H. Randall. Using de bruijn sequences to index a 1 in a computer word. Available on the Internet from http://supertech.csail.mit.edu/papers.html, 1998.

H. Li and R. Durbin. Fast and accurate short read alignment with Burrows-Wheeler transform. Bioinformatics, 25:1754–1760, 2009.

R. Li, C. Yu, Y. Li, T. W. Lam, S.-M. Yiu, K. Kristiansen, and J. W. 0004. SOAP2: an improved ultrafast tool for short read alignment. Bioinformatics, 25(15):1966– 1967, 2009.

Z. Matei, B. W. J., C. Kristal, F. Armando, P. David, S. Scott, S. Ion, K. R. M., and S. Taylor. Faster and more accurate sequence alignment with snap. eprint arXiv, 2011.

G. Myers. A fast bit-vector algorithm for approximate string matching based on dynamic programming. J. ACM, 46(3):395– 415, 1999. ISSN 0004-5411. doi: 10.1145/316542.316550. URL http://doi.acm.org/10.1145/316542.316550.

G. Navarro. A guided tour to approximate string matching. ACM computing surveys (CSUR), 33(1):31–88, 2001.

B. Needleman and C. D. Wunsch. A general method applicable to the search for similarities in the amino acid sequence of two proteins. Journal of Molecular Biology, 1970.

F. Smith and M. S. Waterman. Identification of common molecular subsequences. Journal of Molecular Biology, 147:195–195, 1981.

E. Ukkonen. Algorithms for approximate string matching. Information and control, 64(1-3):100–118, 1985.

H. Xin, D. Lee, F. Hormozdiari, S. Yedkar, O. Mutlu, and C. Alkan. Accelerating read mapping with fasthash. BMC genomics, 14(1):S13, 2013.

